# Molecular portraits of cell cycle checkpoint kinases in cancer evolution, progression, and treatment responsiveness

**DOI:** 10.1101/2020.10.29.361352

**Authors:** E Oropeza, S Seker, S Carrel, A Mazumder, A Jimenez, SN VandenHeuvel, DA Noltensmeyer, NB Punturi, JT Lei, B Lim, S Raghavan, MN Bainbridge, S Haricharan

## Abstract

Cell cycle dysregulation is prerequisite for cancer formation. However, it is unknown whether the mode of dysregulation affects disease characteristics. Here, we conduct comprehensive analyses of cell cycle checkpoint dysregulation events in breast cancer using patient data complemented by experimental investigations in multiple model systems: genetically-engineered mice, patient-derived xenografts, biomatrices, and cell lines. We find that *ATM* mutation predisposes the diagnosis of primary estrogen receptor (ER)+/human epidermal growth factor (HER)2- cancer in older women. Conversely, CHK2 dysregulation induces formation of metastatic, premenopausal ER+/HER2- breast cancer (p=0.001) that is treatment-resistant (HR=6.15, p=0.01). Lastly, while mutations in *ATR* alone are rare, *ATR/TP53* co-mutation is 12-fold enriched over expected in ER+/HER2- disease (p=0.002) and associates with metastatic progression (HR=2.01, p=0.006). Concordantly, ATR dysregulation induces metastatic phenotypes in *TP53* mutant, but not wild-type, cells. These results newly identify a role for distinct cell cycle dysregulation events in determining cancer subtype, metastatic potential, and treatment responsiveness.

**Statement of Significance:** These findings reframe the paradigm of cancer classification by demonstrating that cell cycle dysregulation decisions during malignant transformation can causally direct the type of cancer that evolves, its metastatic potential, and treatment responsiveness. These results provide rationale for delineating mode of checkpoint kinase dysregulation to improve diagnostic and therapeutic choices.

## Introduction

ATM/CHK2 and ATR/CHK1 are cell cycle checkpoint complexes activated by DNA damage^1,2^. Although roles for these proteins in regulating cell cycle progression are complex and often redundant, they can be generalized as ATM/CHK2 and ATR/CHK1 inhibit the cell cycle at G1/S and G2/M phases, respectively^1,3,4^. In cases where prolonged cell cycle arrest is insufficient for DNA repair, these kinases trigger cell death through p53-dependent and - independent mechanisms^5–8^. Consequently, these cell cycle checkpoint kinases are important tumor suppressors across cancer types.

Breast cancer is one of the most frequently diagnosed cancers globally and therefore, one of the most common causes of cancer-related death^9^. Estrogen receptor (ER) status of breast cancer stratifies diagnoses as ER positive (ER+) and negative (ER-)^10^. ER+ breast cancer is far more common than ER-, and predicts response to endocrine therapies that inhibit ER signaling^11,12^. However, ∼20% of ER+ breast cancer patients are intrinsically resistant to endocrine therapy and ∼40% of ER+ breast cancer patients acquire resistance over time^11,13,14^. Triple-negative breast cancer (TNBC) which is characterized by a lack of significant expression of ER, its downstream effector progesterone receptor (PR), and human epidermal growth factor (HER)2 has few available targeted therapeutics, is highly metastatic, and associates with poor patient outcome^10^. Understanding molecular contributions to the evolution of treatment-resistant ER+/HER2- and TNBC can therefore contribute significantly to improving diagnostics and therapeutics for patients.

The importance of cell cycle dysregulation for breast cancer evolution is well established^15,16^. Large and multiple independent epidemiological studies demonstrate the association of germline variants in *ATM* and *CHEK2* with incidence of ER+/HER2- breast cancer^17–21^. Somatic failure to activate ATM/CHK2, through loss of upstream DNA repair signaling for instance, in ER+/HER2- breast cancer can also causally induce resistance to standard endocrine therapies^22–24^. Other studies demonstrate significant associations between high levels of phosphor-ATM and heightened responsiveness to endocrine therapy^25–27^. To date, however, there is a lack of understanding of whether early dysregulation of ATM/CHK2 predisposes the formation of breast cancer that is treatment resistant or aggressively metastatic. Even less is known about the association of *ATR/CHEK1* mutations, either germline or somatic, with breast cancer incidence or outcome. There is evidence that germline mutations in *ATR* are enriched in familial breast cancer patients and that ATR/CHK1 may serve as therapeutic targets in TNBCs^28–30^. But whether mutations in *ATR*/*CHEK1* contribute to the evolution of specific subtypes of breast cancer or to disease progression remains uncertain^31,32^. Overall, the utility of cell cycle checkpoint kinase dysregulation as prognostic markers of disease severity or as predictors of treatment response remains undefined. Previous studies in these areas have suffered from conflicting or inconclusive results largely due to a lack of adequate sample size in patient datasets, and an absence of experimental validation^21,33,34^.

Cell cycle checkpoint dysregulation occurs early in tumor evolution, and there are several ways of achieving this end. Understanding whether specific cell cycle dysregulation events determine the evolution and clinical outcome of different breast cancer subtypes is critical for identifying the potential of each cell cycle protein as prognostic/predictive biomarkers and even as therapeutic targets^21,35^. Here, using breast cancer as a model, we undertake systematic evaluation of the relative contribution of dysregulation of each cell cycle checkpoint kinase to the formation of tumors of distinct subtypes, metastatic progression and treatment responsiveness using a range of informatic and experimental approaches as described below.

## Results

### 1. Mutation of specific cell cycle checkpoint kinase genes promotes the evolution of distinct breast cancer subtypes

Using a meta-dataset comprised of six independent studies (**Figure 1, Figure S1**), we compared the frequency of mutations, both germline and somatic, in each of the four cell cycle checkpoint kinase genes *ATM, CHEK2, ATR*, and *CHEK1* in ER+/HER2- and TNBC samples. ER-/HER2+ samples were excluded from analyses because of insufficient sample size. We included known cancer drivers *ESR1* and *TP53* as positive controls for mutational frequency in ER+/HER2- and TNBC, respectively. As expected, *ESR1* mutations are more common, and *TP53* mutations are less common, in ER+/HER2- than in TNBC tumors (p<1.3e-15 each) (**Figure 1A**). The cumulative frequency of mutation incidence in all four cell cycle kinase genes is comparable between ER+/HER2- and TNBC samples (**Figure 1A**). On an individual gene level, we observed >5-fold enrichment for mutations in *CHEK2* (29/3382 vs 0/640, p=0.001 after adjustment for multiple comparisons) in ER+/HER2- breast cancer samples relative to TNBCs, but no statistically significant enrichment for *ATM, ATR*, or *CHEK1* mutations in either subtype (**Figure 1B**). *CHEK1* mutation is extremely rare in both ER+/HER2- and TNBC and was, therefore, excluded from further analyses. Although mutation of *ATR* alone is not enriched in either ER+/HER2- or TNBC, we found 2-fold enrichment for co-mutation of *ATR* and *TP53* in ER+/HER2- breast cancer (36% *ATR*/*TP53* co-mutated vs 18% *TP53* mutation alone, p=0.002) (**Figure 1C**). Results from this overview analysis indicate that CHK2 is the only cell cycle checkpoint kinase that robustly correlates with the evolution of a specific breast cancer subtype, i.e., ER+/HER2-, when mutated.

**Figure 1.**
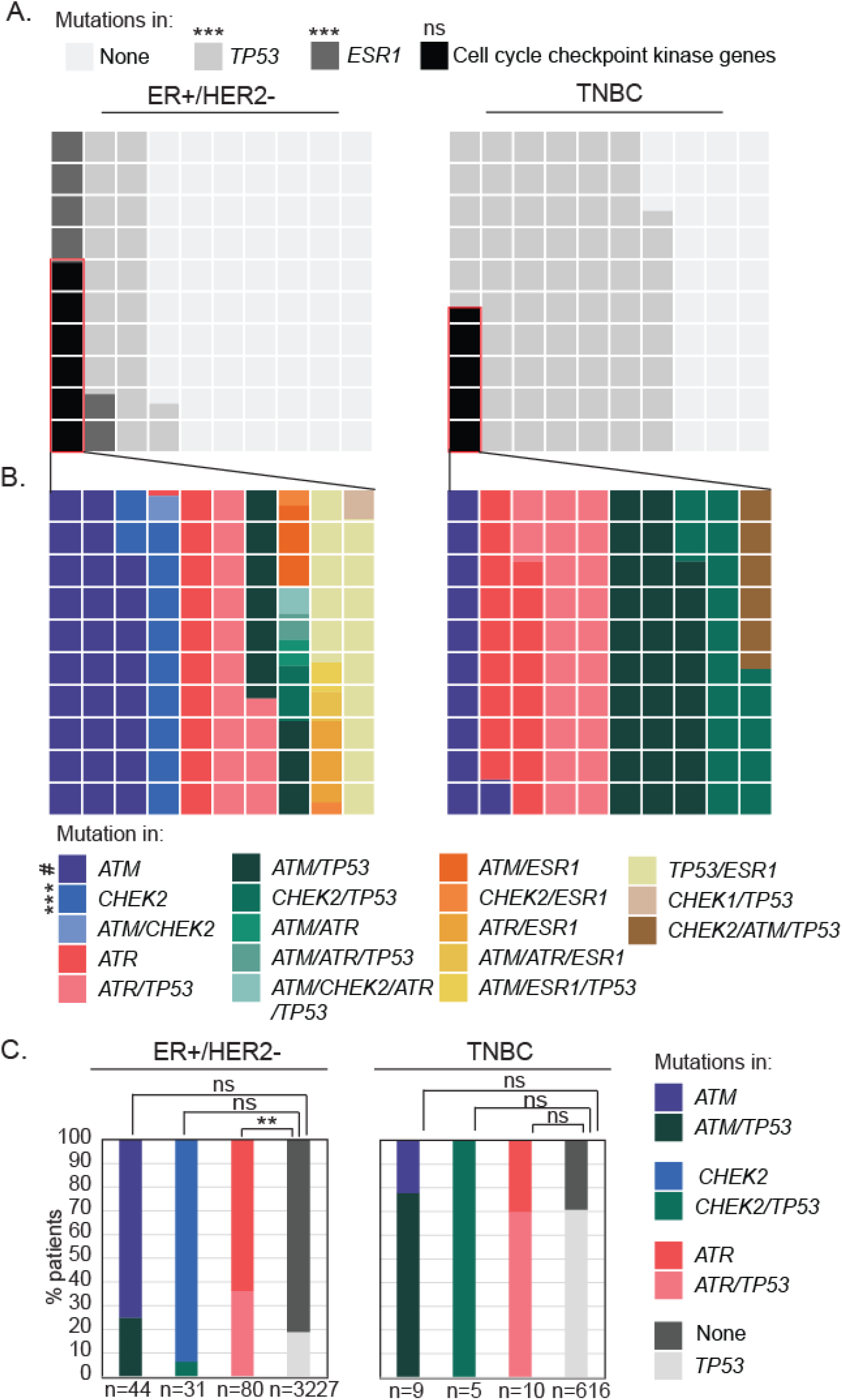
Mutational frequency of cell cycle checkpoint kinase genes differs across breast cancer subtype. **(A-B)** Waffle charts showing *ESR1, TP53* (controls), and cell cycle checkpoint kinase gene (*ATM, ATR, CHEK1*, and *CHEK2*) mutational frequencies in estrogen receptor (ER)+/HER2- vs triple negative breast cancer (TNBC). Each square represents 1%of mutational frequency of specified genes. **(C)** Stacked column graphs representing incidence of mutations in *ATM*/*ATR*/*CHEK2* with and without mutations in *TP53* in the indicated breast cancer subtypes. Fisher’s Exact test determined p-values, and p-values were adjusted for multiple comparison using the Holm’s method. Composition of datasets presented in **Figure S1**. p≤0.1^#^, p≤0.01**, p≤0.001***, ns = not significant.

#### 1.1 CHEK2 mutation enriches for the diagnosis of premenopausal ER+/HER2- breast cancer

The predisposition towards developing ER+/HER2- breast cancer from cells with *CHEK2* mutations indicated the potential involvement of germline variants. As previously established^17^, *ATM* and *CHEK2* are mutated in the germline while *ATR* is not (**Figure 2A**). In this analysis, we found that *CHEK2* is the only cell cycle checkpoint kinase gene that is more likely to be mutated in the germline than somatically (3-fold enrichment in ER+/HER2- breast cancer relative to *ATM* and >50-fold enrichment relative to *ATR*, p=3.7e-12) (**Figure 2A**). We compared the landscape of germline and somatic *CHEK2* mutations and found 3-fold enrichment for deleterious (nonsense, frameshift, or splice site) mutations in the germline group (12/18 vs 4/18 somatic, p=0.02) but no such enrichment among *ATM* mutations (p=0.13) (**Figure 2B**). We next tested whether germline mutations in *CHEK2* alter PR positivity, since ER+/HER2- tumors can be either PR+ (strongly driven by ER signaling) or PR-constituting two distinct breast cancer subtypes: Luminal A and B, respectively^10^. We found that neither germline nor somatic mutations in *CHEK2* impact PR positivity, with tumors remaining predominantly PR+ as is the case with wildtype *CHEK2* tumors (**Figure 2C**). Conversely, we observed that somatic mutations in *ATM* associate with 2-fold enrichment for PR negativity relative to either *ATM* wildtype or germline mutated tumors (p=0.002) (**Figure 2C**).

**Figure 2.**
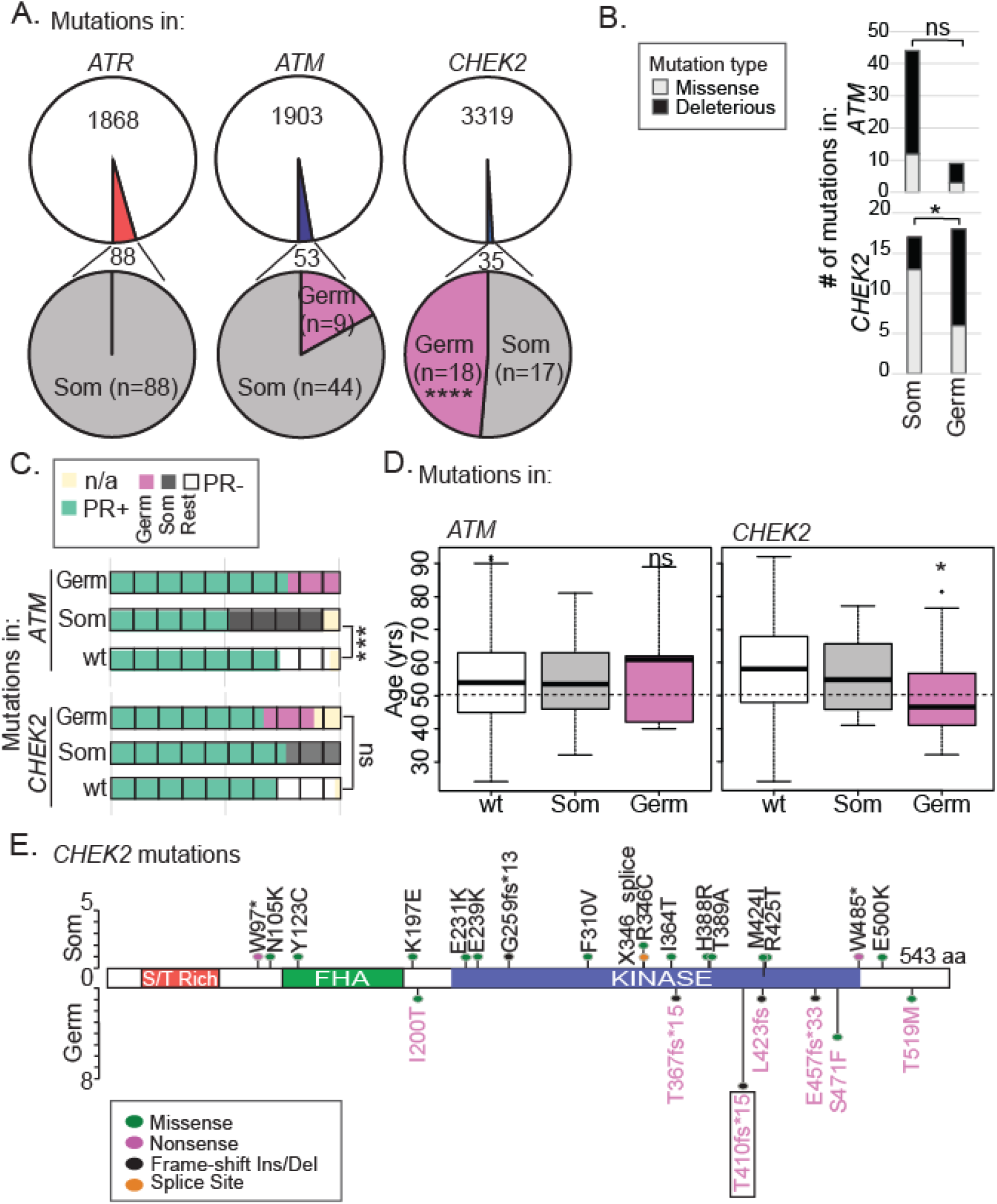
Germline mutations in *CHEK2 drive* associations with younger age at diagnosis of highly PR positive, ER+/HER2- breast cancer. **(A)** Pie charts showing proportion of *ATR, ATM* and *CHEK2* mutations that are somatic and germline in ER+/HER2- breast cancer patients. **(B)** Stacked columns demonstrating percentage of *CHEK2* and *ATM* mutations that are missense and deleterious (frameshift, nonsense or splice site) based on their somatic (Som) or germline (Germ) status in ER+/HER2- breast cancer samples. **(C)** Waffle chart depicting percent ER+/HER2- breast cancer samples with indicated mutations that are progesterone receptor (PR) positive (n/a; PR status not available). **(D)** Box plot indicating median age at diagnosis for indicated groups of breast cancer patients. Error bars describe standard deviation. Dotted line indicates average age at menopause for women in the US. **(E)** Lolliplot of all observed mutations in *CHEK2* labeled in pink if germline and black if somatic. Box indicates the mutation studied experimentally in subsequent analyses. aa, amino acid. Fisher’s Exact Test **(A-C)** and two-tailed independent sample Student’s T-test **(D)** determined p-values. ns, not significant; p≤0.05*, p≤0.01**, p≤0.001***. wt delineates patients with no mutations in genes of interest.

Lastly, we assessed the association of somatic and germline mutations in *CHEK2* and *ATM* with age at diagnosis. We found that germline, but not somatic, mutations in *CHEK2* associate strongly with diagnosis of premenopausal ER+/HER2- disease: median age for *CHEK2* germline carriers is 46 years while that of *CHEK2* somatic or wildtype patients is 55 and 58 years, respectively (p=0.02) (**Figure 2D**).

Conversely, mutations in *ATM* do not impact age at diagnosis: patients with germline or somatic *ATM* mutations remain postmenopausal with median ages of 61 and 54 years, respectively, as expected for ER+/HER2- breast cancer patients (**Figure 2D**). Overall, these data suggest that germline mutations in *CHEK2* contribute to the evolution of ER-driven, premenopausal ER+/HER2- breast cancer.

#### 1.2 CHEK2 mutation induces the evolution of ER+ premalignant growth in a genetically engineered mouse model

To experimentally test whether germline *CHEK2* variants promote the evolution of ER+ cancer in the premenopausal breast, we used a genetically engineered mouse model expressing the *CHEK2*\*1100delC variant (p. Thr410fs*15)^36^, the most common variant in our study (**Figure 2E**). Using this mouse model, we investigated the impact of CHK2 loss on the process of tumor evolution in young and old mice, and after experimental induction of menopause (**Figure 3A**). In mammary gland whole mounts from premenopausal, 5 month old C57/B6 female mice, we identified the formation of macroscopic, preneoplastic mammary lesions in 100% of mice homozygous for the mutant allele (n=5/5), 56% of mice heterozygous for the mutation (n=5/9), and 0% of mice who were wildtype (n=0/5) (p=0.008) (**Figure 3B**).

**Figure 3.**
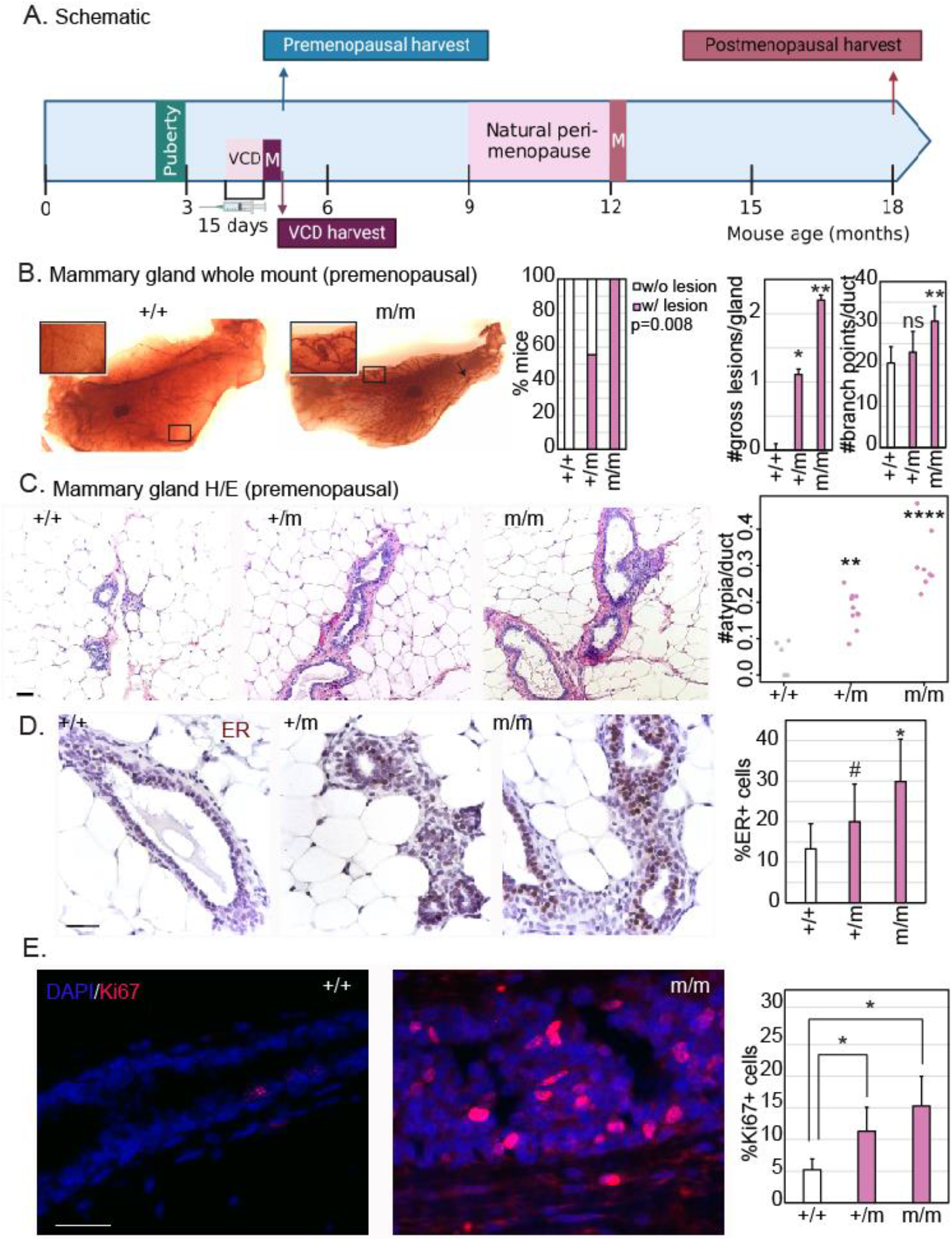
*CHEK2* mutation induces the formation of atypical ER+, highly proliferative mammary lesions in genetically engineered mice. **(A)** Schematic of experimental design. VCD, 4-vinylcyclohexene diepoxide; M, menopause. **(B)** Representative images of whole mounted mammary glands (1.5x) with cleared fat pads showing mammary ductal structure which was used to quantify the incidence (w/, with; w/o, without) and number of gross (macroscopic) mammary lesions (representative 6x magnification shown in inset) and the number of branch points in mammary ducts, represented in bar graphs. Statistical differences in incidence of lesions were tested using Fisher’s Exact test, and in number of lesions and branch points using a two-tailed Student’s T-test. **(C-E)** Representative images and accompanying bar graph quantification of the number of microscopic atypia using hematoxylin and eosin (H/E) staining **(B)**, percent atypical cells that are ER positive by immunohistochemistry **(C)**, and percent proliferating cells using immunofluorescence for Ki67 **(D)**. Scale bars = 20um. Two-tailed Student’s T-test derived all p-values. For all panels, wildtype (+/+), heterozygous (+/m) and homozygous (m/m) *CHEK2*\*1100delC mice were harvested at 20 weeks (5 months) of age. Error bars in all bar graphs represent standard deviation. ER, estrogen receptor; ns, not significant; p≤0.1^#^, p≤0.05*, p≤0.01**, p≤0.001***. Associated data validating results in the FVB background are presented in **Figure S2**.

Homozygous mutant mice also have multiple macroscopic lesions per mammary gland (p=0.0004) and heavy side branching (**Figure 3B**), indicators of unchecked proliferation, although ductal length is unperturbed compared to wildtype mice (**Figure S2A**). These findings are supported by the observed increase in microscopic, atypical lesions in the mammary ducts of homozygous (6-fold increase over wildtype, p=1.7e-5) and heterozygous (4-fold increased over wildtype, p=0.001) mutant mice (**Figure 3C**). Preneoplastic lesions in homozygous mutant mice are also heavily ER+ with a 2-fold higher rate of median ER positivity than wildtype ducts (p=0.02) (**Figure 3D**) and highly proliferative (**Figure 3E**). Since mouse strain backgrounds can significantly influence mammary phenotypes, we confirmed these phenotypes in an FVB background, reproducing the increase in macroscopic preneoplastic lesions (p=0.04) (**Figure S2B**), microscopic atypia (p=0.002) (**Figure S2C**), ER positivity (p=0.003) (**Figure S2D**) and proliferation (p=0.01) (**Figure S2E**) in homozygous mutants relative to wildtype controls.

#### 1.3 Mutant CHEK2 induced premalignant cancer evolution is suppressed in the postmenopausal mammary epithelium

To investigate whether mutant *CHEK2*-induced premalignant growth is altered by menopausal status, we first compared premenopausal (5 month old) and postmenopausal (18 month old) mammary glands from heterozygous and homozygous *CHEK2*\*1100delC mutant female mice. We found that the mammary epithelia of 18 month old mutant mice harbor significantly fewer macroscopic lesions (**Figure 4A, Figure S3A**) than their 5 month old counterparts, suggesting that the postmenopausal mammary environment suppresses premalignant evolution.

**Figure 4.**
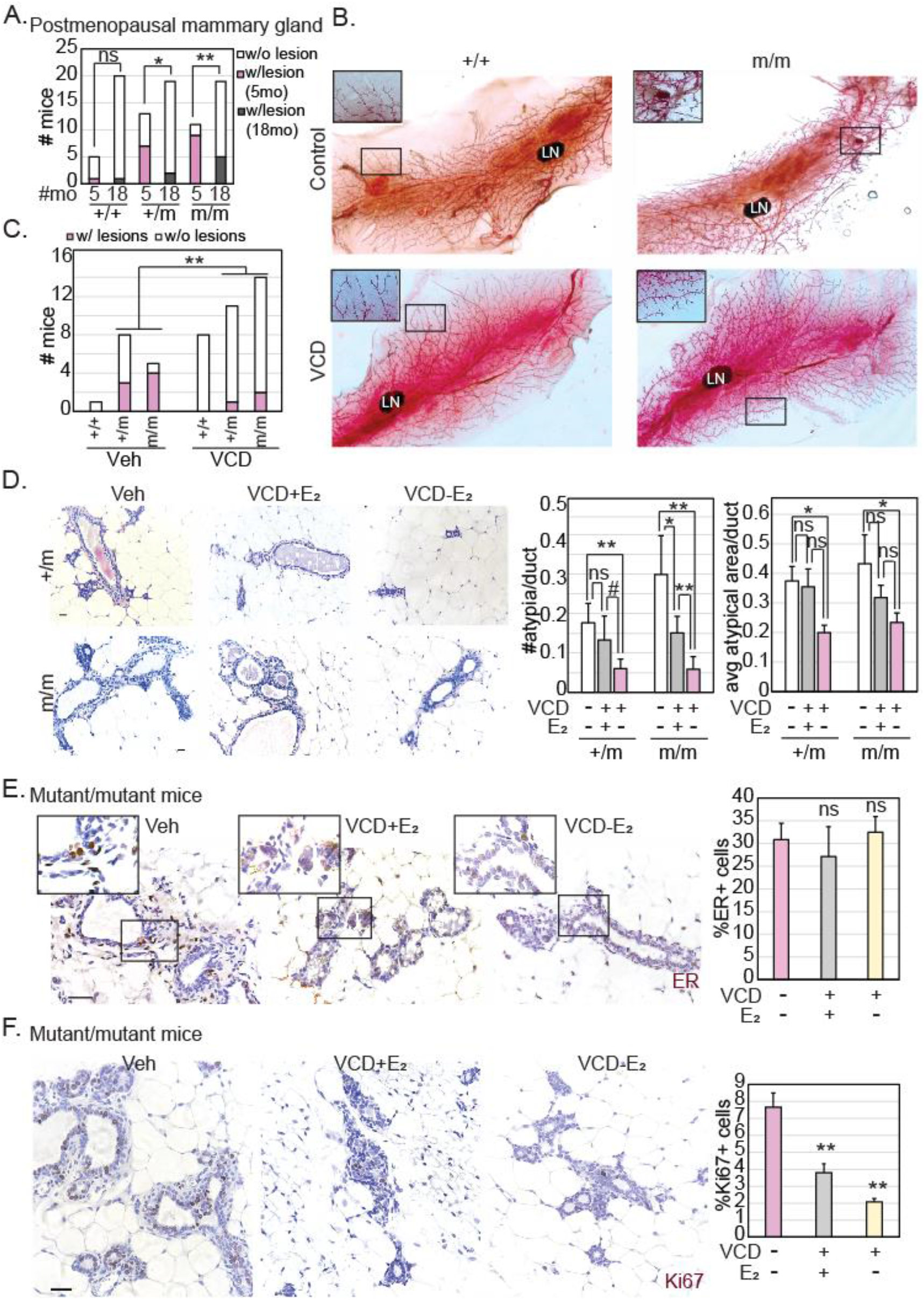
*CHEK2* mutation induces the formation of highly proliferative mammary lesions preferentially in premenopausal mice. **(A)** Stacked column graph representing the proportion of wildtype (+/+), heterozygous (+/m) and homozygous (m/m) *CHEK2*\*1100delC mutant female mice at 5 (premenopausal) and 18 (postmenopausal) months of age. **(B-C)** Representative images **(B)** of whole mounted mammary glands (1.5x; inset magnification 2.5x) from mice from each specified genotype, treated with or without 4-vinylcyclohexane diepoxide (VCD), with cleared fat pads showing mammary ductal structure which was used to quantify the incidence (w/, with; w/o, without) of gross (macroscopic) mammary lesions (representative magnification shown in inset), represented as a stacked column graph **(C)**. Statistical differences in incidence of lesions tested using Fisher’s Exact test. LN, Lymph Node. **(D-F)** Representative images and accompanying bar graph quantification of the number of microscopic atypia using hematoxylin and eosin (H/E) staining **(D)**, percent atypical cells that are estrogen receptor (ER) positive by immunohistochemistry **(E)**; inset magnified 2x), and percent proliferating cells using immunofluorescence for Ki67 **(F)** under the specified treatment conditions (Vehicle, Veh; 4-vinylcyclohexane diepoxide, VCD; E_2_, beta-estradiol). Scale bars = 20μm. Two-tailed Student’s T-test determined p-values. Mice were harvested at 20 weeks (5 months) of age for **B-F**. Whiskers in all bar graphs represent standard deviation. ns, not significant; p≤0.1^#^, p≤0.05*, p≤0.01**, p≤0.001***. Associated data presented in **Figure S3**.

As a more direct test of the impact of the postmenopausal mammary environment on *CHEK2* mutation-induced breast cancer evolution, we used the 4-vinylcyclohexene diepoxide (VCD) model^37^ to induce menopause in heterozygous and homozygous mutant, female mice. Mice were either administered VCD or placebo injections for 15 days and sacrificed 6 weeks after administration, at 5 months of age (schematic in **Figure 3A**). As expected, we observed almost complete suppression of serum estradiol levels within 4 weeks of administration of VCD (p=0.002) (**Figure S3B**). Importantly, we observed macroscopic preneoplastic mammary lesions in 80% of the vehicle-treated homozygous mutant mice (4/5) and 40% of heterozygous mice (3/8), compared to only 14% (2/14) and 9% (1/11) of VCD-treated controls (cumulative p=0.005) (**Figure 4B-C**). The decrease in preneoplastic incidence is echoed at the microscopic level with 75% and 85% decrease in lesion incidence in heterozygous and homozygous mutants respectively (p=0.001) (**Figure 4D**). Existing atypia in VCD-treated mutant mice also demonstrate significant shrinkage in area (**Figure 4D**). The number of ER+ cells in atypia in VCD-treated mice remains comparable to that in vehicle treated controls although levels of ER appear lower (**Figure 4E**). Irrespective of ER positivity, proliferation in preneoplastic cells is significantly suppressed in VCD-treated homozygous mutants compared to vehicle-treated controls (**Figure 4F**).

To test whether the suppression of preneoplastic evolution in postmenopausal *CHEK2* mutant mammary glands is due to loss of mitogenic estrogen stimuli caused by suppression of ovarian function after menopause, we included a control group of mice whose drinking water was supplemented with estradiol during and after administration of VCD. We found that estrogen supplementation partially rescues the preneoplastic phenotype in VCD-treated mice (**Figure 4D**). However, lesions arising in estrogen supplemented VCD-treated mice have a distinct morphology, unlike that seen in vehicle-treated premenopausal mice (**Fig 4D**) indicating that the observed rescue of macroscopic lesions might be an independent effect of estrogen stimulation of mammary cells. In support of this hypothesis, estrogen supplementation does not prevent the proliferative inhibition caused by VCD-induced menopause (p=0.0005) (**Figure 4F**). Together, these data suggest a causal role for CHK2 loss in inducing premenopausal ER+ breast cancer evolution that is at least partially independent of estrogen-mediated mitogenesis.

### 2 Individual cell cycle checkpoint kinase genes have distinct impact on disease progression when mutated

We next tested whether mutations in cell cycle checkpoint kinase genes modulate metastatic progression in breast cancer patients. As expected, in our breast cancer meta-dataset, incidence of *ESR1* mutations is highly enriched, and of *TP53* moderately enriched, in metastatic ER+/HER2- breast cancer compared to primary^38,39^ (**Figure 5A**). Amongst cell cycle checkpoint kinase genes, we observed 1.5-fold enrichment for *ATM* mutation (alone, not in combination with any other gene of interest) in primary ER+/HER2- breast cancer relative to metastatic disease (p=0.003) (**Figure 5B**). Strikingly, *ATM* is the only cell cycle checkpoint kinase gene enriched for mutation in the primary setting, as both *CHEK2* (p=0.001, **Figure 5C**) and *ATR* mutations (p=0.0002, **Figure 6A**) associate with metastatic disease.

**Figure 5.**
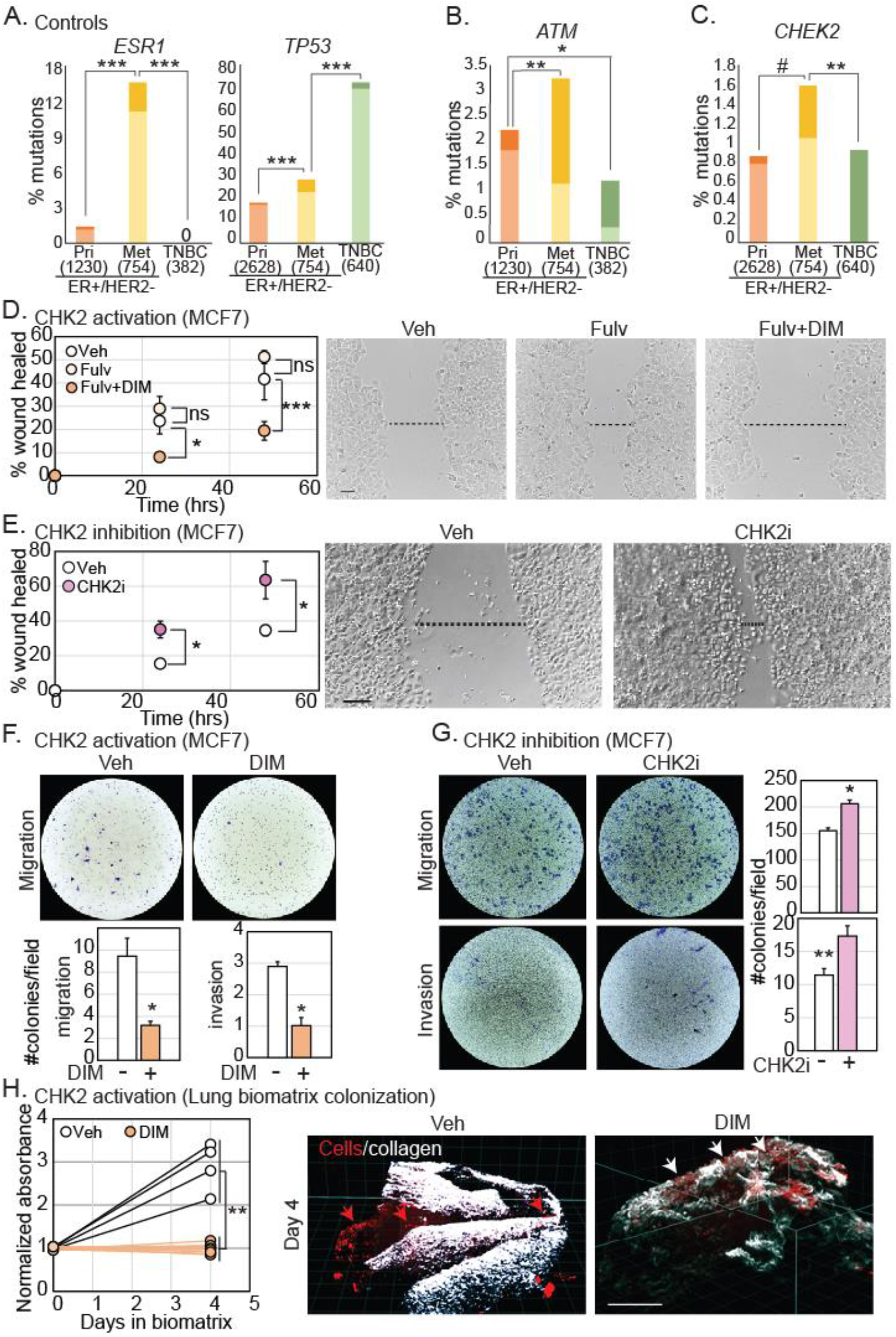
CHK2 loss promotes metastatic ER+/HER2- disease. **(A-C)** Bar graphs representing frequency of mutations in *ESR1* and *TP53* **(A**, control genes), *ATM* **(B)**, and *CHEK2* **(C)** in estrogen receptor (ER)+/HER2-primary (Pri) versus metastatic (Met) breast cancer and in triple negative breast cancer (TNBC). Darker shading indicates incidence of mutations in multiple genes of interest while lighter shading indicates mutation of only the specified gene of interest. P-values were derived by comparing the incidence of light shaded columns between the three breast cancer subtypes in a Fisher’s Exact test. Holm’s adjustment for multiple comparisons was conducted. Sample sizes in parentheses. **(D-E)** Representative images of wound healing assay of ER+/HER2- breast cancer cell line, MCF7, treated with vehicle (Veh), 100 nM Fulvestrant (Fulv) or the combination of 10μM di-indolyl methane (DIM), a CHK2 activator and fulvestrant **(D)**, or 100nM CHK2 inhibitor dihydrate (CHK2i) **(E)** at 48 hours. Dot plots representing quantification of area of scratch at 0, 24 and 48 hours with error bars depicting standard deviation. **(F-G)** Representative images of transwell migration (top) and invasion (bottom) assays at 48 hours after treatment with vehicle, DIM or CHK2i, along with bar graphs depicting quantification. DIM treatments were done in media with charcoal stripped serum + 10 pM beta-estradiol., while CHK2i assays were done in media with full serum. Error bars represent standard deviation. **(H)** Dot plots quantifying cell clusters that invade into the bioengineered lung matrix (collagen) and establish micrometastatic colonies along with representative images showing RFP-tagged T47D cell clusters and the collagen matrix (white). Micrometastatic colonies established after invasion into the collagen matrix are indicated by red arrows and cell clusters that remain above the collagen unable to invade into the matrix are indicated by white arrows. Scale bars=200 μm. Independent sample Student’s T-test determined p-values for **D-H**. not significant, ns; p≤0.1^#^; p≤0.05*; p≤0.01**; p≤0.001***. Supporting data validating activity of compounds used and independent confirmation in a second cell line in **Figure S4**.

**Figure 6.**
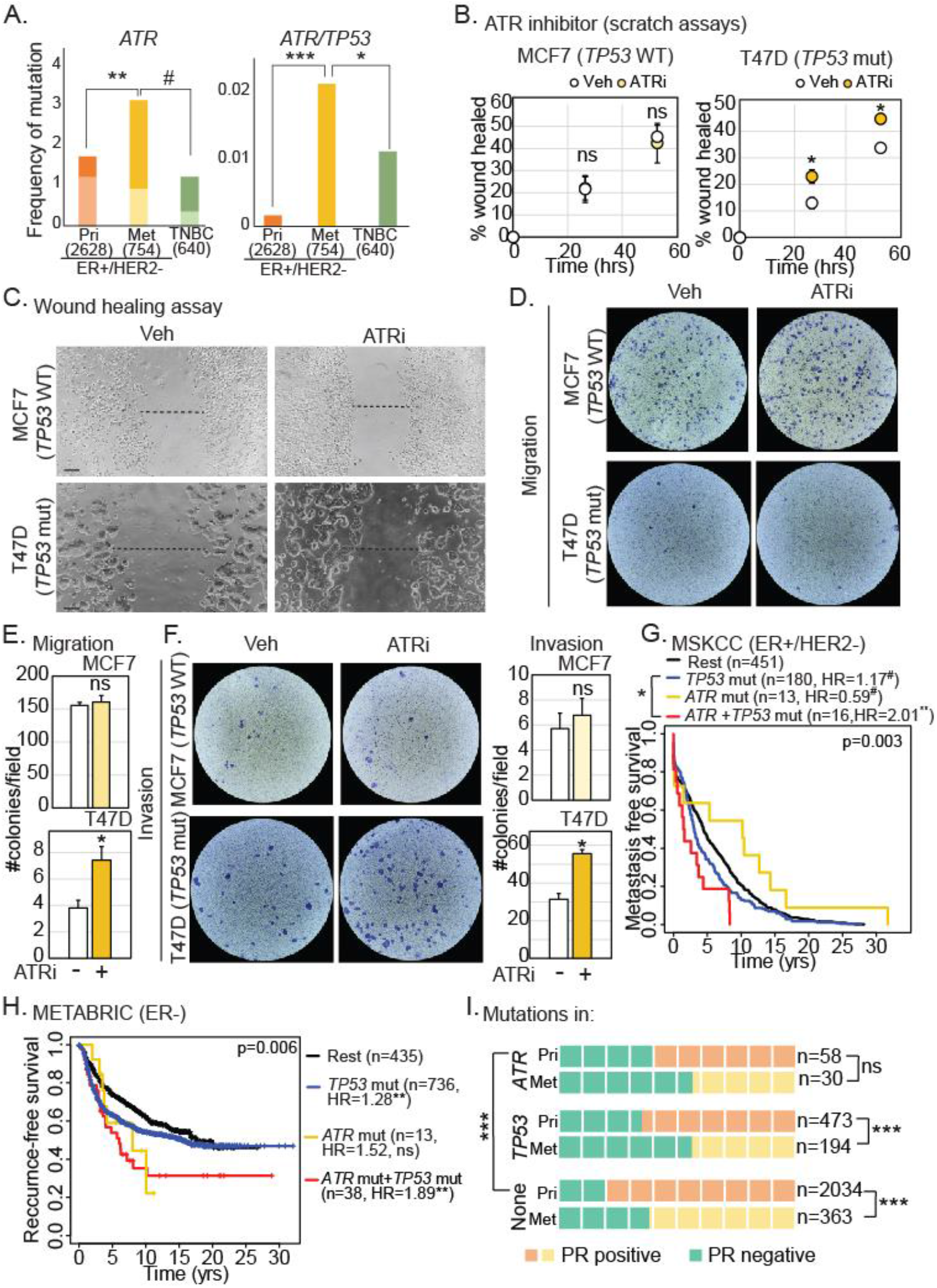
ATR dysregulation promotes metastatic breast cancer dependent on *TP53* status. **(A)** Bar graphs representing frequency of *ATR* and *TP53* mutations in estrogen receptor (ER)+/HER2- primary (Pri) versus metastatic (Met) breast cancer and in triple negative breast cancer (TNBC). The darker shading indicates that there were mutations in multiple genes of interest while the lighter shading indicates that only the specified gene of interest was mutated in those samples. P-values were derived by comparing the light shaded columns between the three breast cancer subtypes. Sample sizes in parentheses. **(B-C)** Representative images **(C)** of wound healing assay of ER+/HER2- breast cancer cell lines, MCF7 (*TP53* wildtype; wt) and T47D (*TP53* mutant; mut), treated with vehicle (Veh) or ATR inhibitor (ATRi) at 48 hours. Dot plots representing quantification of area of scratch at 0, 24 and 48 hours with error bars depicting standard deviation **(B). (D-F)** Representative images of transwell migration **(D)** and invasion **(F)** assays at 48 hours after treatment with vehicle or ATRi, along with bar graphs depicting quantification **(E, F)**. Error bars represent standard deviation. **(G-H)** Kaplan-Meier survival curves representing metastasis-free **(G)** and local+distant recurrence-free **(H)** survival in the specified genotypic cohorts. HR, hazard ratio. **(I)** Waffle chart depicting percent ER+/HER2- primary and metastatic breast cancer samples with indicated mutations that are progesterone receptor (PR) positive. Fisher’s Exact test determined p-values for **A** and **I** (with Holm’s adjustment for multiple comparisons), two-tailed Student’s T-test for **B-F**, and log rank test for **G-H**. not significant, ns; p≤0.1^#^; p≤0.05*; p≤0.01**; p≤0.001**; in none of the genes of interest, none. Supporting data validating activity of compounds used and independent confirmation in other patient tumor datasets in **Figure S5**.

#### 2.1 CHK2 dysregulation causally impacts metastasis in breast cancer cells

To understand the functional relevance of CHK2 dysregulation in promoting the metastasis of ER+/HER2- cancer, we experimentally dysregulated CHK2 in two independent cell lines (MCF7: **Figure 5** and T47D: **Figure S4**). First, we activated CHK2 exogenously using fulvestrant, a standard endocrine therapy that induces modest CHK2 activation (**Figure S4A-B**), and di-indolyl methane (DIM), a robust, small molecule CHK2 activator^40,41^ (**Figure S4A-B**). In wound healing assays, treatment with endocrine therapy alone is not sufficient to impact cell migration but administration of DIM in addition to endocrine therapy significantly inhibits motility (**Figure 5D**: MCF7 (19% vs 42% wound healed, p=0.008), **Figure S4C**: T47D (6% vs 32%, p=0.01)). Conversely, inhibition of CHK2 using a small molecule inhibitor, CHK2-inhibitor dihydrate^42^ significantly promotes motility in wound healing assays in both cell lines (**Figure 5E**: MCF7 (63% vs 45% wound healed, p=0.01), **Figure S4D**: T47D (50% vs 33%, p=0.02)). These observations were orthogonally replicated through assessment of migration and invasion in transwell assays in both MCF7 and T47D cells (**Figure S4E-F**).

To rule out the effect of endocrine treatment in the role of CHK2 activation on motility and invasiveness, wound healing, transwell migration, and invasion assays were also conducted with either DIM or CHK2 inhibitor treatment alone, without addition of endocrine therapies. Results from these experiments confirmed the role of CHK2 activation in inhibiting migration and invasion (**Figure 5F-G, Figure S4G**) and of CHK2 inhibition in promoting these phenotypes (**Figure 5F-G, Figure S4H**), respectively in each of these assays.

Lastly, we seeded RFP-ragged T47D spheroids (**Figure S4I**) into bioengineered porcine lung biomatrix scaffolds. We visualized and quantified the ability of these cells to invade and colonize the lung matrix and establish cell clusters representing micrometastatic colonies. We observed establishment of micrometastatic colonies within four days in vehicle-treated T47D cells, while this ability is completely abrogated by treatment with DIM (**Figure 5H;** 2.8-fold increase in absorbance relative to day 0 in control vs no increase in DIM-treated cells, p=0.006). Importantly, the DIM treatment did not significantly affect spheroid growth outside the biomatrix (**Figure S4I**), suggesting a selective impact of CHK2 activation on the ability of cells to invade and establish themselves at distant organs.

#### 2.2 ATR dysregulation causally impacts metastasis in TP53 mutant breast cancer cells

Unlike mutations in *ATM* or *CHEK2*, mutations in *ATR* alone are uncommon in both primary and metastatic ER+/HER2- breast cancer. However, co-incident mutation of *ATR* and *TP53* is 12-fold enriched in metastatic ER+/HER2- breast cancer over primary disease (p=0.002), and 2-fold enriched over TNBC (p=0.02) (**Figure 6A**). Additionally, in a pan-cancer analysis of TCGA and MSKCC data, *ATR* is the only cell cycle checkpoint kinase that is co-mutated with *TP53* (q<0.001, log_2_ odds ratio=1.23). *ATM* mutation is mutually exclusive with *TP53* mutation (q=0.01, log_2_ odds ratio=-0.28) and *CHEK2* mutation is not correlated with *TP53* mutation (q=0.1, log_2_ odds ratio=0.48).

In experimental assays, administration of a validated small molecule ATR inhibitor^30^ (by Western blotting and immunofluorescence; **Figure S5A**) significantly promotes motility in wound healing (**Figure 6B-C**) assays, and migration (**Figure 6D-E**) and invasion (**Figure 6F**) in transwell assays, in *TP53* mutant T47D cells. However, ATR inhibition has no effect on motility or invasion of *TP53* wildtype MCF7 cells (**Figure 6B-F**). These observations are supported by analyses of metastatic progression in ER+/HER2- breast cancer patients: *ATR* mutation alone does not associate with metastasis-free survival, but *ATR*/*TP53* co-mutation associates significantly with worse metastasis-free survival in two independent datasets (**Figure 6G, Figure S5B**).

This co-mutation of *ATR/TP53* is also an independent prognostic factor for increased metastatic recurrence in ER+/HER2- breast cancer patients (**Figure S5C**). Given that *TP53* mutation is a hallmark of ER- breast cancer and *ATR*/*TP53* co-mutation appears to be a pan-cancer feature, we also tested the association of *ATR*/*TP53* co-mutation with metastasis-free survival in ER- disease. We found that *ATR*/*TP53* co-mutation associates significantly with poor metastasis-free survival in ER- breast cancer in both datasets analyzed (**Figure 6H, Figure S5D**). In accordance with these data, while primary ER+/HER2- breast cancer is normally highly PR positive with decreasing levels in metastatic disease, mutations in *ATR* and *TP53* associate with 1.7-fold higher likelihood of PR negativity even in primary disease relative to wildtype tumors (p=2.7e-10) (**Figure 6I**).

Together, these data support distinctive roles for individual cell cycle checkpoint kinases in metastatic progression of breast cancer, with dysregulation of *ATM* associating with primary ER+/HER2- disease, *CHEK2* with metastatic ER+/HER2- breast cancer, and *ATR* with metastatic disease that is ER agnostic but reliant on *TP53* mutation.

#### 2.3 CHK2 dysregulation causally impacts treatment responsiveness in breast cancer

We next investigated the association of mutations in cell cycle checkpoint kinase genes with response to endocrine treatment using two independent datasets, MSKCC^43^ and METABRIC^44^. We found that *CHEK2* mutations in metastatic ER+/HER2- breast cancer patients, whether germline or somatic, associate with shorter progression-free survival (average of 183 days) on frontline endocrine therapy relative to patients with *CHEK2* wildtype disease (584 days) (MSKCC, p=0.03) (**Figure S6A**). We found a similar association between incidence of germline *CHEK2* mutations and worse relapse-free survival in primary ER+/HER2- breast cancer patients in METABRIC (HR=6.15, p=0.01) (**Figure 7A**). This association remains significant in a proportional hazards assessment including known prognostic factors of PR status, tumor stage, age at diagnosis and type of administered endocrine therapy (**Figure S6B**) and is not detectable in primary ER+/HER2- breast cancer patients who did not receive endocrine therapy (**Figure S6C**). Of note, we did not observe an association between somatic *CHEK2* mutations and poor outcomes in METABRIC. This is likely because is no significant enrichment for deleteriousness in germline vs somatic *CHEK2* mutations in metastatic breast cancer samples in MSKCC. In contrast, this enrichment is detectable in METABRIC (6/6 germline mutations are deleterious vs 3/12 somatic mutations, p=0.009 by Fisher’s Exact test). We confirmed the association between CHK2 dysregulation and poor outcome for ER+/HER2- breast cancer patients using proteomic data for phosphor-CHK2 (TCGA: HR=2.0, p=0.02) (**Figure 7B**) and validated the observed enrichment for CHK2 dysregulation in ER+/HER2- disease relative to either HER2+ (1.5-fold) or TNBC (10-fold) at the protein level (**Figure 7C**, p=0.0004).

**Figure 7.**
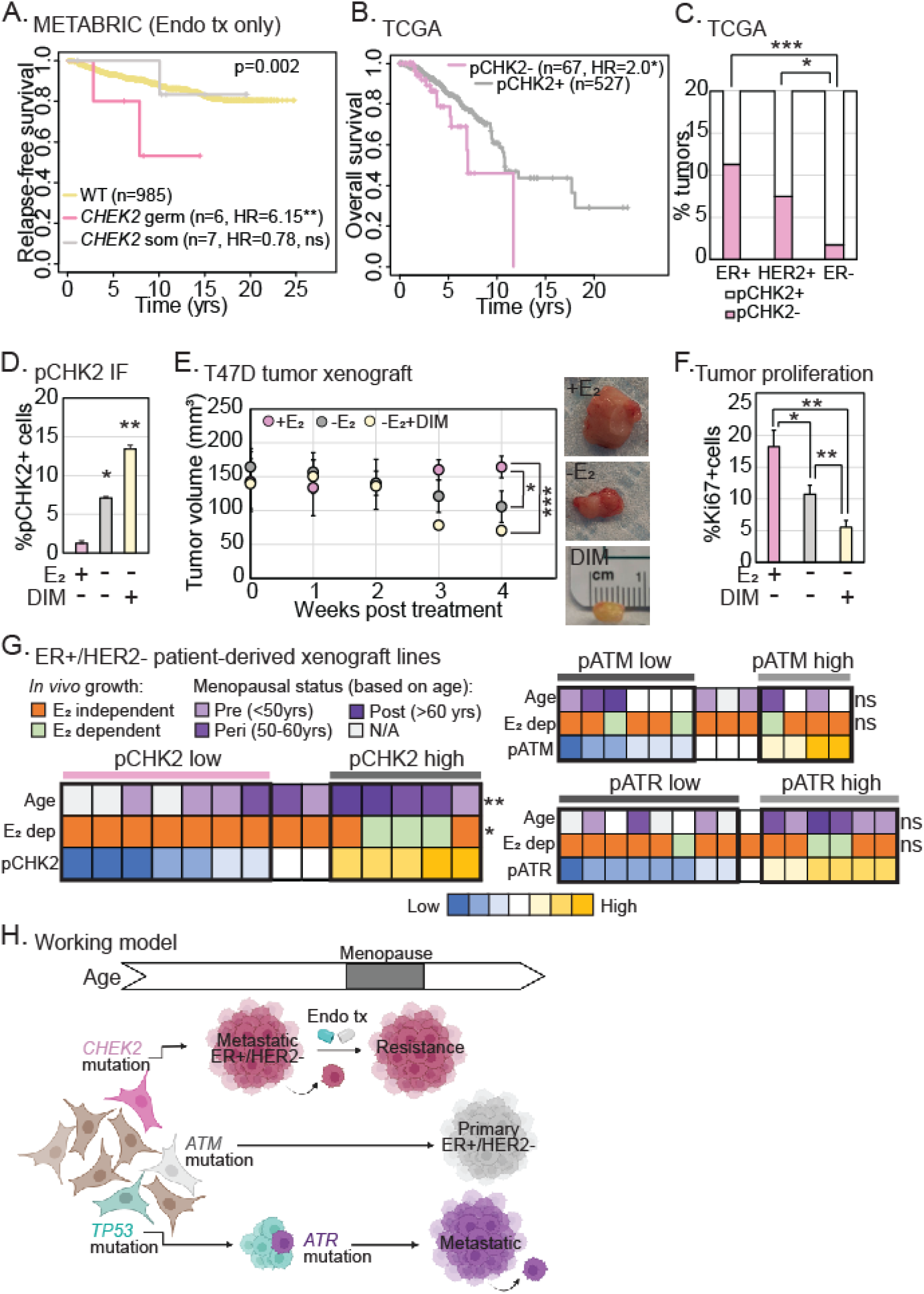
CHK2 activation promotes responsiveness to standard endocrine therapy. **(A-B)** Kaplan-Meier survival curves measuring the specified outcomes in patients with mutations in **(A)** or downregulation of **(B)** CHK2 relative to those proficient for CHK2 (wildtype, WT). Log rank test determined p-values. Supporting data for proportional hazards assessment and validation in independent datasets are presented in **Figure S6**. HR, hazard ratios. **(C)** Stacked column graph representing incidence of CHK2 downregulation as measured by reverse phase proteomics array in ER+/HER2- breast cancer samples from TCGA. Fisher’s Exact test determined p-values. **(D, F)** Bar graphs representing quantification of percentage of pCHK2+ cells and proliferating cells (Ki67+) in xenograft tumors derived from ER+/HER2- breast cancer cell line, T47D, grown in mice with estradiol supplementation in drinking water (Veh), estrogen deprivation (endocrine therapy, Endo) and CHK2 activator, di-indolyl methane (DIM) incorporated in their chow (treatment, tx). Two-tailed Student’s T-test determined p-values. Error bars represent standard deviation. **(E)** Dot plot depicting mean tumor volumes in mice xenografted with T47D cells and treated with specified treatments. Two-tailed Student’s T-test determined p-values. Three mice/group. Representative images of tumors from specified treatment conditions. **(G)** Heat maps indicating protein levels of pCHK2, pATM, and pATR across a panel of ER+/HER2- breast cancer patient-derived xenografts. Ability of tumors to grow in the absence of estrogen supplementation (E_2_) is represented as well as age at which the patient was diagnosed with breast cancer. Low (mean-standard deviation) and high (mean+standard deviation) phospho/total protein levels of cell cycle checkpoint kinases were compared for statistically significant differences in age at diagnosis and ability to grow in the absence of estrogen using Fisher’s Exact test. **(H)** Working model depicting the impact of mutations in different cell cycle checkpoint kinase genes on the type of breast cancer a patient may develop and clinical consequences in terms of treatment response and metastatic recurrence. ns, not significant; p≤0.1^#^, p≤0.05*, p≤0.01**, p≤0.001***. Associated data presented in **Figure S6**.

To experimentally validate a link between CHK2 dysregulation and endocrine therapy resistance, we administered DIM to mice bearing xenografted tumors from ER+/HER2- T47D breast cancer cells. Estrogen deprivation alone activates CHK2 (**Figure 7D**) and inhibits tumor growth (p=0.04) (**Figure 7E**). However, the addition of DIM activates CHK2 twice as much as estrogen deprivation alone (p=0.02) (**Figure 7D**; 11-fold increase over estrogen-supplemented tumors, p=0.007) to further suppress tumor growth (p=0.005) (**Figure 7E**) and proliferation (p=0.008) (**Figure 7F**). These *in vivo* data support the hypothesis that CHK2 activation mediates responsiveness to endocrine therapies.

As further experimental validation, we conducted a proteo-genomic analysis of phosphor-ATM, phosphor-ATR and phosphor-CHK2 levels across a panel of ER+/HER2- PDX lines. We found that low levels of CHK2 phosphorylation predict estrogen-independent growth across PDX lines *in vivo* (**Figure 7G**; 7/7 phosphor-CHK2 low PDX lines are estrogen-independent compared to 2/5 phosphor-CHK2 high lines, p=0.04). We also found that ER+/HER2- breast cancer PDXs with low levels of phosphor-CHK2 are more likely to be derived from premenopausal breast cancer patients (**Figure 7G**; median age for pCHK2 low PDXs is 38.5 vs 57 in pCHK2 high lines, p=0.009) in accordance with data from the GEMM and the mutational analysis of germline *CHEK2* variants in patient data presented above. Of note, we found no such correlations with either ATM or ATR phosphorylation (**Figure 7G**).

Overall, these data provide some of the first evidence for divergences in the causal impact of ATM, CHK2 and ATR inactivation on the type of breast cancer a patient develops and on disease progression, i.e., metastatic potential and responsiveness to endocrine therapies (**Figure 7H**).

## Discussion

While cell cycle checkpoint kinases are well recognized as tumor suppressors across cancer types, and the efficacy of inhibitors of these proteins (primarily ATR and CHK1) in targeting cancer has been frequently investigated^45^, we lack a systematic understanding of the relative contributions of each of these kinases to cancer initiation and progression. The findings presented in this work constitute the first systematic analysis of the implications of dysregulation of each of these kinases on tumor subtype formation and disease progression that incorporates patient tumor data analysis with experimental validation of associations, using breast cancer as a model.

We find that women who are carriers of deleterious, germline variants in *CHEK2* are predisposed to the incidence of premenopausal ER+/HER2- breast cancer. Our experimental demonstration in genetically engineered mice that *CHEK2* mutation-induced tumorigenesis requires the premenopausal mammary environment is important, initial evidence of a complex interplay between hormones, aging, and cell cycle checkpoint signaling. This finding is supported by previous reports of a 4% incidence rate for *CHEK2*\*1100delC mutations in premenopausal breast cancer patients^46^ versus a 0.7% incidence rate in postmenopausal breast cancer patents^47^ in a population background of ∼0.2% (calculated from ClinVar). These data raise the hypothesis that different cell cycle checkpoint kinases may be preferentially involved in promoting cancer in young people, which requires in-depth investigation. These results also raise the translationally important question^48^ of whether breast cancer screening in *CHEK2* germline variant careers should be modulated based on age and menopausal status to prevent overdiagnosis and overtreatment.

Additionally, we find that breast cancer patients with either somatic or germline mutations in *CHEK2* are more likely to be diagnosed with metastatic than primary ER+/HER2- breast cancer. These data are supported by previous studies showing increased incidence of germline *CHEK2* mutations in metastatic breast cancer relative to primary disease^49^, and worse recurrence-free survival outcomes for *CHEK2* mutation carriers^50^. To our knowledge, the results presented here constitute the first experimental validation of a direct role for CHK2 inhibition in promoting metastatic phenotypes in ER+/HER2- breast cancer cells, although studies from another group indicate a role for CHK2 activation in epithelial mesenchymal transition^51,52^, often a precursor to metastasis. Mechanisms that underlie this causal association warrant further study.

We also find that breast cancer patients with *CHEK2* mutations appear resistant to endocrine monotherapy, which targets the ER signaling pathway, despite their cancer being highly PR positive. High PR positivity is considered a sign of dependence on ER signaling. In cell line and patient-derived xenograft tumor growth studies, we demonstrate that CHK2 dysregulation causally alters estrogen dependence *in vivo*. The efficacy of a small molecular CHK2 activator in inhibiting tumor growth *in vivo* also suggests new therapeutic avenues for next generation cell cycle-based cancer therapeutics. Since ATM and CHK2 inhibitors are largely ineffective in cancer clinical trials^53^, development of activators of these kinases might afford more complete cell cycle control especially in conjunction with CDK4/6 inhibitors that have shown efficacy in the clinic^54^. Overall, these findings provide a comprehensive portrait of the distinctive impact of CHK2 dysregulation on the evolution of premenopausal, metastatic, highly PR positive, ER+/HER2- breast cancer that is likely to resist standard endocrine monotherapies.

Conversely, we find that *ATM* mutations enrich for the incidence of primary ER+/HER2- breast cancer that is preferentially PR-negative. PR negativity in ER+/HER2- breast cancer patients associates with more aggressive disease that may be less responsive to endocrine therapies^21,55^. However, we did not find associations between *ATM* mutations and poor patient outcomes in this study. While ATM and CHK2 are often paired together in canonical cell cycle regulation and DNA damage response^25,56,57^, the results of our study suggest that they have distinct cellular functions that may explain their individual impact on breast cancer presentation and progression. Previous reports of distinct roles for these two kinases in G1 checkpoint regulation may constitute one mechanism underlying their differential impact on cancer phenotypes^58^. Larger datasets and more mechanistic studies are required to comprehensively assess the impact of ATM dysregulation on disease progression and patient outcome.

Finally, as a novel observation, we uncover a role for co-mutation of *ATR* and *TP53* in promoting breast cancer metastasis. The ability of p53 mutational context to modulate the oncogenic/tumor suppressor potential of many driver genes (e.g. Myc^59^, TLR4^60^) is well-recognized. In the same vein, both in patient tumor data and in experimental analyses, we find that loss of ATR only impacts tumor phenotypes and biology in the context of dysregulated p53. Further, the association of this co-mutation with metastasis-free survival appears significantly stronger than that of mutations in either gene alone. This dichotomy may explain why previous epidemiological studies considering tumors of both wildtype and mutant p53 status failed to find associations between ATR dysregulation and breast cancer outcome. Moreover, these results are translationally critical in informing the use of ATR inhibitors in the clinic to selectively target *TP53* wildtype cancers. It is also important to point out that this co-mutation effect is only observed for *ATR* and not for the other cell cycle checkpoint kinases. In fact, in a pan-cancer analysis of TCGA data, *ATR* is the only cell cycle checkpoint kinase co-mutated with *TP53*.

Overall, this systematic analysis of the association of individual cell cycle checkpoint kinase genes with clinically relevant tumor characteristics and breast cancer patient outcome suggests that the mode of cell cycle dysregulation during tumorigenesis may be leveraged to improve the characterization of cancer subtypes. While breast cancer is an excellent cancer type for these proof-of-concept analyses, the implications of these findings for other cancer types warrant investigation. With successful clinical implementation of CDK inhibitors for certain types of cancer, there is renewed interest in understanding how different cyclin and CDK dependencies in cancer cells can guide therapeutic decisions^61^. As upstream regulators of these cyclins and CDKs, ATM, ATR, CHK1 and CHK2, are amongst the most common dysregulation events that promote cancer susceptibility across cancer types^62^. The results of this study, by demonstrating clear divergences in the impact of dysregulation of ATM, ATR, and CHK2 on important tumor characteristics, shed new insight into how early decisions to turn off cell cycle regulation can direct the course of ensuing disease. Simple inhibition of each kinase as tested in the clinic^63,64^ may, therefore, not be an optimal solution. Further evidence that the early loss of specific cell cycle checkpoint kinases could essentially serve as a decision point for the evolution of different cancer subtypes may argue for an improved system of cancer classification based on the mode of cell cycle checkpoint inactivation to guide the selection of therapeutics. Development of such prognostic and predictive stratifiers could provide new strategies to match CDK inhibitors or next generation cell cycle checkpoint activators to the individual cell cycle dependencies of each patient’s tumor.

## Notes

### Author contributions

Data collection, curation and analysis (EO, SP, NP, AM, SvH, DN, JTL, BL, SR, MNB, SH). Conceptual oversight (MNB, SH). Preparing manuscript (EO, SP, SC, NP, SH). Writing and editing of manuscript (EO, SP, SC, AM, BL, SR, MNB, SH).

### Data availability

The datasets were derived from sources in the public domain: TCGA : Cancer Genome Atlas Network, at https://www.cancer.gov/about-nci/organization/ccg/research/structural-genomics/tcga, cBioPortal [https://www.cbioportal.org/study/summary?id=brca_tcga_pub]

METABRIC : cBioPortal [https://www.cbioportal.org/study/summary?id=brca_metabric]

MSKCC : DOI. https://doi.org/10.1016/j.ccell.2018.08.008, Cancer Cell 2018

BROAD : cBioPortal [https://www.cbioportal.org/study/summary?id=brca_broad]

MBCP : The Metastatic Breast Cancer Project [ww.mbcproject.org], cBioPortal [https://www.cbioportal.org/study/summary?id=brca_mbcproject_wagle_2017]

### Citation diversity statement

It is established across academic fields that papers authored by scientists who are white and Asian men are over-cited compared to those authored by scientists from URGs (including women). In an effort to address any citation biases that may exist in our paper, we have compiled the gender and race of each first and last author cited. We were then able to determine the proportion of papers authored by scientists from URGs (including women) compared to those authored by scientists who identify as white or Asian men. In this article, 56.1% of the papers cited were authored by at least one woman in a senior position (either first of last), 15.8% were authored by at least one non-white author in a senior position, and 10.5% were undetermined.

## Methods

### Datasets

Six datasets with mutational data from primary and metastatic breast cancer patients were combined for initial analyses (see **Figure1** and **Figure S1**). The results included here include the use of data from The Metastatic Breast Cancer Project (https://www.mbcproject.org/),a project of Count Me In (https://joincountmein.org/). Analyses regarding patient outcomes were conducted in two independent datasets: MSKCC^65^ and METABRIC. Details of each dataset in **Supplementary Methods**.

### Mutational analysis

All protein changing mutations in *TP53, ESR1, ATM, CHEK2, ATR* and *CHEK1* genes were included irrespective of category (i.e., missense, nonsense, etc) or predicted pathogenicity. Protein changing mutations in *TP53* and *ESR1* genes served as controls as they are known drivers of TNBC and ER+/HER2- breast cancer, respectively. Mutational frequency was calculated based on total number of mutations divided by patient count.

### Tumor characteristics

Tumor PR status and age of diagnosis served as categorical variables to determine associations between incidence of mutations and patient/tumor characteristics. Fisher’s exact test was used to determine p-values by comparing different categories such as ER+/HER2- vs TNBC or germline vs somatic status for PR positivity while a two-tailed Student’s T-test was used for continuous age differences.

### Genetically-engineered Mice

The mice in the 5-month premenopausal experiment were from either strain C57BL/6N (line: Atm1BrdChek2tm1a(EUCOMM)Hmgu/JMmucd) which were received from MMRRC (Cat#047089-UCD) or strain 129/Sv*BlackSwiss*FVB/N (line: Chek2tm1Pjs/Mmnc) which were also received from MMRRC (Cat#01411-UNC). The mice in the 18-month postmenopausal and VCD-induced menopause experiment were only from strain 129/Sv*BlackSwiss*FVB/N. The 129/Sv*BlackSwiss*FVB/N lines was backcrossed to FVB mice for 6 generations to stabilize the strain background. For all mice, genotyping was done when mice were 4-6 weeks old. The C57BL/6N mice were genotyped according to MMRRC protocol with bands expected around 384 bp, with the 129/Sv*BlackSwiss*FVB/N genotyped similarly with bands expected around 500 bp.

### *In vivo* experiments

For the 5-month and 18-month experiments, mice were genotyped at 4-6 weeks and housed in random groups between 6-8 weeks old, then harvested at 5 months and 18 months, respectively. Mice were palpated once monthly until tumors were palpable. Once a tumor was palpable, palpations were done weekly. None of the 5-month mice developed palpable tumors. Of the 59 mice in the 18-month experiment, 23 mice (11 +/+, 5 +/m, 7 m/m) were found dead with cause of death unknown. Details of the VCD-induced menopause experiment (as per published protocol^66^) are in **Supplementary Methods**. All mammary fat pads were harvested from the 5-month premenopausal, 5-month induced menopause, and 18-month postmenopausal experiments. The left #4 and left #2/3 were fixed in 4% paraformaldehyde and paraffin embedded, the right #2/3 was snap frozen, and right #4 was whole-mounted and stained with neutral red as previously described^55^. Mice for the xenograft experiment were 6-8 week old NOD/SCID mice (from Sanford Burnham Prebys). Mice were injected with T47D cells suspended in Matrigel (Corning Cat# 356234), and randomized into 3 treatment groups (+E_2_, - E_2_, -E_2_+DIM) when tumors reached 100mm in diameter. Estradiol was supplemented into sterile deionized water at a concentration of 8 ug/mL, and DIM (MedChemExpress) was given in diet form (Research Diets). Tumors were injected into the left #4, measured twice weekly, and harvested as previously described^22^.

### Immunostaining

Immunofluorescence was performed based on manufacturer’s instructions and as per previously published protocols^22^. Antibodies used include phosphor-Chk2 (Cell Signaling Technology Cat#2197S), Estrogen Receptor and Ki67 (Abcam Cat#ab16667).

Immunohistochemistry was performed based on manufacturer’s instructions. Sections were first deparaffinized, then endogenous peroxidases were quenched using 3% H202, and antigen retrieval was done using 1x citric acid buffer. The blocking buffer for Estrogen receptor (EMD Millipore Cat#04820MI, 1:200) and Ki67 (Abcam Cat#ab16667, 1:200) staining was 2% goat serumPrimary antibodies were left overnight in 4 degrees and followed by anti-rabbit secondary (Vector Laboratories Cat# BA-1000). Next, sections were incubated in avidin-biotin complex solution (Vector Laboratories Cat#PK-6100), stained with peroxidase substrate (Vector Laboratories Cat#SK-4800), and counterstained in hematoxylin. Images were captured on an Echo Revolve microscope.

### Migration and invasion assays

Wound healing and transwell migration assays were performed to assess metastatic potential. For wound healing assays, 2.5×10^5^ T47D cells were plated in 6-well plates and incubated for 24 hours. Using a 20μL pipette tip, a scratch was made in the center of the plate and pictures were taken at 0, 24 and 48 hours. Fresh media with 100 nmol/L of ATR inhibitor (cat# S8007; Selleckchem), CHK2 inhibitor (cat# C3742; Sigma-Aldrich) or 1mmol/L of CHK2 activator, di-indolyl methane (DIM) (cat# sc-204624B, Santa Cruz Biotechnology) were added and incubated for 48 hrs. Pictures were taken at 48 hrs to quantify wound healing. For transwell migration and invasion assays, 2.0×10^5^ cells in 200 μL of media (no FBS + ATR or CHK2 inhibitors) were added in transwell inserts (Cat# 353182; Falcon). For invasion assay, inserts were coated with Matrigel (cat# 356234; Corning) in 1:3 dilution with media without FBS. Inserts were placed on 12 well plate with 750 μL of cell culture media. Fixation and staining were carried out after overnight incubation. Transwells were placed in fixative (cat# 22-122911; Fisher Scientific) followed by 5 mins staining in Solution (cat# 22-122911; Fisher Scientific). DIM transwell experiments were conducted in media with charcoal stripped serum and estradiol supplementation, while all inhibitor experiments were conducted in media with full serum. Inserts were dried and pictures were taken using Echo microscope.

### Biomatrix assays

The T47D breast cancer cell line was cultured on a hanging drop array to form 3D tumor spheroids over the course of 4 days^67,68^. On day 3, the spheroids were either treated with 10 μM DIM inhibitor or fed with fresh media for control. After compact spheroid formation was confirmed (day 4), the cells were seeded onto a decellularized lung biomatrix scaffold to establish engineered breast cancer lung metastasis (BCLM) per previously established protocols^69^. The cells were allowed 2 hours for matrix attachment before undergoing MTS metabolic assay or being submerged in culture medium (+/- DIM) for continued growth. Following 4 days of culture, engineered BCLM were harvested to measure metabolic activity via MTS assay.

### Western blotting

Western blotting was conducted as previously described. Cells were exposed to 48 hours of ATR or CHK2 inhibitor/activator treatments. All antibodies were diluted in 1x TBST in 1:1000 dilution and incubated overnight at 4 °C. Antibodies used were Phospho-ATR (Cat# 2853; Cell Signaling), ATR (Cat# 2790; Cell Signaling), Phospho-Chk2 (Cat# 2197; Cell Signaling), Chk2 (Cat# 2662; Cell signaling) and GAPDH (Cat# sc-47724; Santa Crutz).

### Survival and disease progression analyses

Outcome measures used were relapse-free, metastasis-free and local/distant recurrence-free survival. Only samples with survival metadata were included. Cox Proportional Hazards calculated hazard ratios and p-values. Tumor stage, PR status, age at diagnosis and classes of endocrine therapy were included in multivariate analyses.

### Statistical analyses

Missing data were imputed with “NA” from mutation and survival data analysis. Samples classifying for more than one category were treated as separate set for statistical comparisons. Independent sample Student’s t-test was used for age comparisons and Fisher’s Exact test was used for categorical data. Log rank test calculated p-values for survival analyses. For every analysis where multiple hypotheses were tested, Holm’s adjustment for multiple comparisons was used.

