## Supplementary figures for "Molecular portraits of cell cycle checkpoint kinases in cancer evolution, progression, and treatment responsiveness"

**Supplementary Figures and Legends**

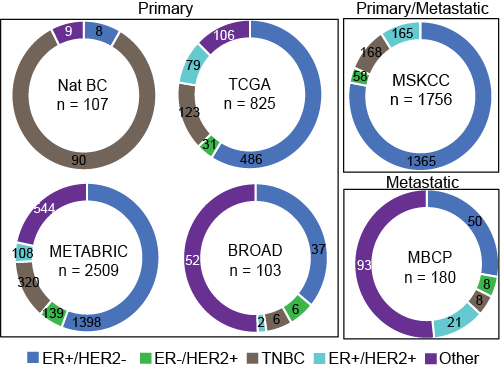

**Supplementary Figure 1: Description of samples by patient datasets.** Donut plots demonstrate the proportion of tumors from each of the six datasets used divided based on hormone receptor (HR) status and HER2 positivity. TNBC, Triple negative breast cancer. Other indicates samples where one or more of the receptor status variables had missing information. Nat BC, Nature British Columbia; TCGA, The Cancer Genome Atlas; MSKCC, Memorial Sloan Kettering Cancer Center; METABRIC, Molecular Taxonomy of Breast Cancer International Consortium; BROAD, Broad Institute; MBCP, Metastatic Breast Cancer Project. Supports analyses presented in **Figure 1**.

**Supplementary Figure 2. *CHEK2* mutation induces the formation of highly proliferative mammary lesions in genetically engineered mice (FVB background). (A)** Bar graph representing quantification of total ductal length (arbitrary units, AU) in mammary glands from female *CHEK2**1100delC mice in the C57/B6 strain background. **(B)** Stacked column graph quantifying incidence (w/lesion, with lesion; w/o lesion, without lesion) and bar graph representing number of gross (macroscopic) mammary lesions. Statistical differences in incidence of lesions tested using Fisher’s Exact test, and in no. of lesions using Student’s T-test. **(C-E)** Representative image for ER immunofluorescence **(D)** and bar graph quantification of the number of microscopic atypia using hematoxylin and eosin staining **(C)**, percent atypical cells that are ER positive by immunohistochemistry **(D)**, and percent proliferating cells using immunofluorescence for Ki67 **(E)**. Scale bars = 20µm. Student’s T-test derived p-values. For all panels, wildtype (+/+), heterozygous (+/m) and homozygous (m/m) *CHEK2**1100delC mice were harvested at 20 weeks (5 months) of age. **(B-E)** represent mice in the FVB strain background. Error bars in all bar graphs represent standard deviation.ER, estrogen receptor; ns, not significant; p≤0.1^#^, p≤0.05*, p≤0.01**, p≤0.001***. Supports data presented in **Figure 3**.

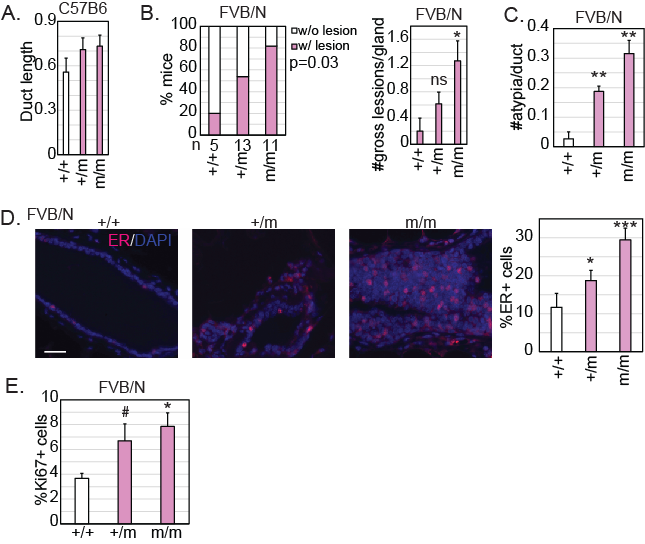

**Supplementary Figure 3. *CHEK2* mutation induces the formation of highly proliferative mammary lesions in young mice. (A)** Representative images and accompanying bar graph quantifying number of macroscopic (gross) lesions per mammary gland in female wildtype (+/+), heterozygous (+/m) and homozygous (m/m) *CHEK2**1100delC mice harvested at 18 months of age. Scale bars = 20µm. Inset magnification 2.5x. Statistical differences in number of lesions were determined using Student’s T-test. Supports data presented in **Figure 4A**. **(B)** Bar graph quantification of serum estradiol levels in female mice treated with control (Ctrl) or VCD (4-vinylcyclohexane diepoxide) injections. Student’s T-test derived p-values. Error bars in all bar graphs represent standard deviation LN, lymph node; ns, not significant; p≤0.1^#^, p≤0.05*, p≤0.01**, p≤0.001***. Supports data presented in **Figure 4B-F**.

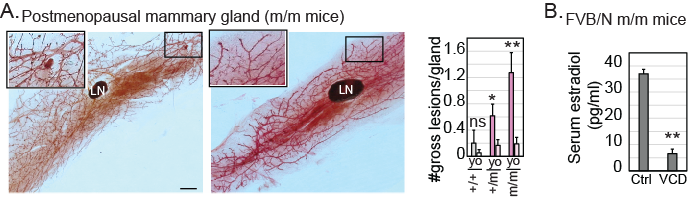

**Supplementary Figure 4: CHK2 dysregulation modulates metastatic phenotypes in ER+/HER2- breast cancer cells. (A)** Bar graphs representing quantification of nuclear phosho-CHK2 positivity in MCF7 cells treated with vehicle (Veh), Fulvestrant (Fulv, 100nM) or DIM (10µM) for 36 hours, along with representative photomicrographs. Scales bars represent 20µm. P-values were derived using a Student’s T-test. **(B)** Western blot demonstrating efficacy of 100nM fulvestrant (Fulv), 10µM DIM (DIM) and 100nM CHK2 inhibitor dihydrate (CHK2i) in altering levels of phosphorylated CHK2 in T47D cells after 36 hours of treatment. **(C-D)** Dot plots representing quantification of area of scratch at 0, 24 and 48 hours after specified treatments with error bars depicting standard deviation. Student’s T-test for p-values. **(E-H)** Bar graphs representing quantification of transwell migration and invasion assays at 48 hours after specified treatments. Error bars represent standard deviation. Student’s T-test derived p-values. All DIM experiments were conducted in media containing charcoal stripped serum supplemented with beta-estradiol while all inhibitor experiments were conducted using media with full serum. not significant, ns; p≤0.1^#^; p≤0.05*; p≤0.01**; p≤0.001***. Supports data presented in **Figure 5**.

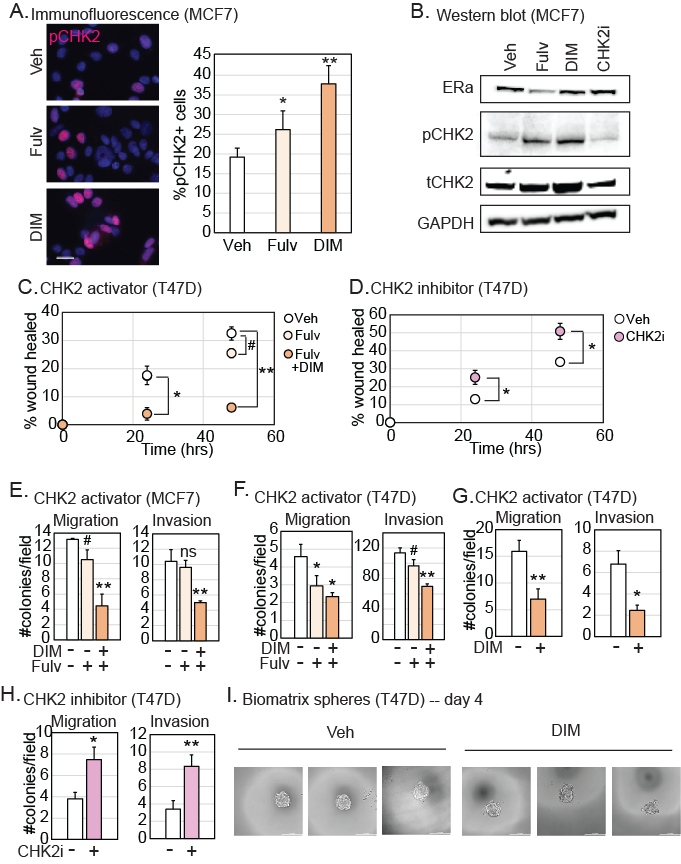

**Supplementary Figure 5. Association of ATR dysregulation with metastasis. (A)** Bar graph with quantification of pATR positivity assayed by immunofluorescence (IF) of MCF7 and T47D cells, and Western blot demonstrating efficacy of ATR inhibitors (100nM for 48 hours) on inhibiting ATR phosphorylation in MCF7 cells. GAPDH used as loading control. **(B&D)** Kaplan-Meier survival curves measuring the specified outcomes in tumors with mutations in specified genes. Log rank test determined p-values. **(C)** Forest plots depicting the hazard ratio of indicated survival parameters in a Cox proportional hazards analysis with standard prognostic factors: Progesterone Receptor (PR), tumor stage, age at diagnosis (dx) and category of endocrine therapy (aromatase inhibitor, AI; tamoxifen, TAM). Boxes indicate the hazard ratio and error bars indicate the 95% confidence intervals. A yellow box indicates a higher hazard ratio, and a blue box indicates a lower hazard ratio relative to reference. Supports data presented in **Figure 6**. ATRi, ATR inhibitor; ns, not significant; HR, hazard ratios;yrs, years; Wt, wildtype; PR, progesterone receptor; ER, estrogen receptor; Unk, unknown.*, p<0.05; **, p<0.01; ***, p<0.001.

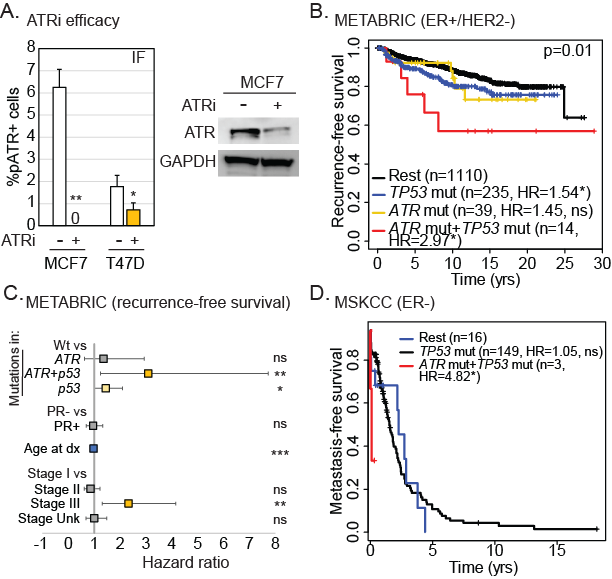

**Supplementary Figure 6. CHK2 loss associates with worse patient outcome. (A)** Stacked column graph depicting the length of time for which metastatic ER+/HER2- breast cancer patients stayed on their first (E1) and second (E2) lines of endocrine treatment before progressing categorized based on the presence of somatic (Som) or germline (Germ) mutations in *ATM* or *CHEK2*. Fisher’s Exact test determined p-values. **(B)** Forest plots depicting the hazard ratio of indicated survival parameters in a Cox proportional hazards analysis with standard prognostic factors: Progesterone Receptor (PR), tumor stage, age at diagnosis (dx) and category of endocrine therapy (aromatase inhibitor, AI; tamoxifen, TAM). Boxes indicate the hazard ratio and error bars indicate the 95% confidence intervals. A yellow box indicates a higher hazard ratio, and a blue box indicates a lower hazard ratio relative to reference. **(C)** Kaplan-Meier survival curves measuring the specified outcomes in patients with mutations in specified cell cycle checkpoint kinase genes. Log rank test determined p-values. HR, hazard ratios; ns, not significant; p≤0.1^#^, p≤0.05*, p≤0.01**, p≤0.001***. Supports data presented in **Figure 7**.

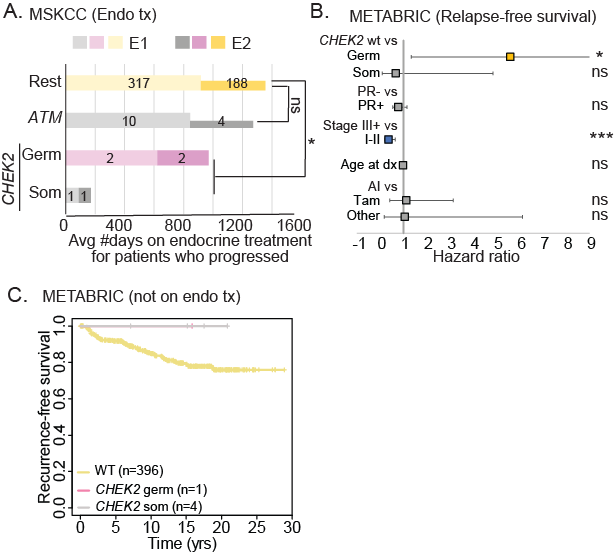

**Supplementary Methods**

Details of datasets

*MSKCC, Cancer Cell 2019* ^14^: Clinical and mutational data (*ESR1, TP53, ATM, ATR, CHEK2* and *CHEK1*) collected from 1756 patients with primary hormone receptor positive (i.e. ER and/or PR+) (HR+)/HER2- (n=1365 ), HR-/HER2+ (n=58), TNBC (n=168) and HR+/HER2+ (n=165) breast cancers.

*TCGA, Nature 2012* ^64^: Downloaded from cBioPortal^65^ for clinical and mutational (*ESR1*,*TP53*, *ATM*, *ATR*, *CHEK2* and *CHEK1*) analysis in December 2019.The data set is composed of clinical and mutational data collected from 825 patients with primary HR+/HER2- (n=486), HR-/HER2+ (n=31), TNBC (n=123), HR+/HER2+ (n=79) and undetermined (n=106) breast cancers.

*METABRIC Nature 2012* ^41^ *& Nature Communication 2016* ^66^*:* Downloaded from cBioPortal for clinical and mutational (*ESR1*, *TP53*, *ATR* and *CHEK2*) analysis in December 2019.The data set is composed of clinical and mutational data collected from 2509 patients with primary HR+/HER2- (n=1398), HR-/HER2+ (n=139), TNBC (n=320), HR+/HER2+ (n=108) and undetermined (n=544) breast cancers.

*Broad, Nature 2012* ^67^: Downloaded from cBioPortal for clinical and mutational (*ESR1, TP53, ATM, ATR, CHEK2* and *CHEK1*) analysis in March 2020.The data set is composed of clinical and mutational data collected from 103 patients with primary HR+/HER2- (n=37), HR-/HER2+ (n=6), TNBC (n=6), HR+/HER2+ (n=2) and undetermined (n=52) breast cancers.

*MBCP, Provisional, February 2020* ^68^: Downloaded from cBioPortal for clinical and mutational (*ESR1, TP53, ATM, ATR, CHEK2* and *CHEK1*) analysis in March 2020.The data set is composed of clinical and mutational data collected from 180 metastatic patients with HR+/HER2- (n=50), HR-/HER2+ (n=8), TNBC (n=8), HR+/HER2+ (n=21) and undetermined (n=93) breast cancers.

*British Columbia, Nature 2012* ^69^*:* Downloaded from cBioPortal for clinical and mutational (*ESR1, TP53, ATM, ATR, CHEK2* and *CHEK1*) analysis in March 2020.The data set is composed of clinical and mutational data collected from 107 patients with primary HR+/HER2- (n=8), TNBC (n=90) and undetermined (n=9) breast cancers.

**VCD-induced menopause** For the VCD experiments, female mice were genotyped at 5 weeks, housed in random groups at 10 weeks, and started on estrogen treatment. VCD (Sigma-Aldrich Cat#S453005) injections began when mice were 12 weeks old and administered IP at a dosage of 160 mg/kg. Control mice were injected with PBS in sesame oil. Mice were harvested at 8 weeks post injection, and serum was collected via cardiac puncture and sent to the Ligand Core at UVA for estradiol analysis.
